# Feed-based delivery of plasmid DNA using milk-derived extracellular vesicle-polyethylenimine complexes enables hepatic transgene expression in goldfish

**DOI:** 10.64898/2026.10.20.760587

**Authors:** Qiwen Zheng, Hiu Mai, Wenliangyu Li, Shuisheng Li, Binbin Zheng

## Abstract

Oral administration through feed offers a non-invasive, cost-effective, and scalable approach for gene delivery in aquaculture. However, gastrointestinal degradation and intestinal barriers severely limit the oral delivery of plasmid DNA (pDNA). In this study, we developed a ternary nanocarrier composed of bovine milk-derived extracellular vesicles (MEVs), polyethylenimine (PEI), and pDNA (MPPs) to investigate its potential for functional gene delivery to the goldfish (*Carassius auratus*) liver (hepatopancreas). The resulting MPPs exhibited a relatively uniform particle size (120.8 ± 12.0 nm), high pDNA loading efficiency, and protection against nuclease-mediated degradation. Formulation optimization in HEK293T cells identified an MEV:PEI:pDNA mass ratio of 16.7:4.1:1, which supported robust EGFP reporter expression. Following oral administration of DiD-labeled MEVs through feed, fluorescence signals were detected in the goldfish hepatopancreas within 2 h and remained detectable at 6 h. After a single feeding of MPP-containing feed, hepatic luciferase mRNA expression peaked at 24 h and gradually declined to near-baseline levels by 72 h. During 14 consecutive days of feeding, luciferase protein expression was detectable from day 3, with progressively increased protein abundance and enzymatic activity observed on days 7 and 14. Collectively, these findings demonstrate the feasibility of feed-based MPP administration for inducing functional transgene expression in the goldfish hepatopancreas, providing a promising foundation for developing non-invasive nucleic acid delivery strategies in aquaculture.

## INTRODUCTION

Plasmid DNA (pDNA)-mediated gene delivery holds considerable promise for aquaculture, with potential applications in disease prevention, growth regulation, and metabolic health management in farmed fish^1,2^. However, translating this potential into large-scale aquaculture practice requires effective and practical administration methods^3^. Although injection provides reliable dosing, it is labor-intensive and technically challenging, particularly in juvenile fish, and inevitably causes handling stress and tissue injury^3^. Immersion-based delivery, meanwhile, is limited by the susceptibility of nucleic acids to degradation in aquatic environments and their poor tissue penetration. In contrast, oral administration through feed offers a non-invasive, cost-effective approach suitable for large-scale application^4,5^. However, orally administered pDNA must withstand gastrointestinal degradation, traverse the intestinal mucus and epithelial barriers, and reach target tissues in a sufficiently intact form to support cellular uptake and nuclear entry^6,7^. These successive barriers severely limit the oral bioavailability of naked pDNA. Therefore, developing carriers that protect pDNA while facilitating intestinal absorption and subsequent tissue delivery is essential for advancing feed-based gene delivery in aquaculture^7,8^.

Among potential target organs, the liver (hepatopancreas) is particularly attractive for gene delivery in fish because of its central roles in nutrient metabolism, growth regulation, and innate immunity^9,10^. Efficient delivery of pDNA to this organ could provide new opportunities for improving growth performance, feed utilization, and hepatic health. Chitosan-and poly(lactic-co-glycolic acid) (PLGA)-based carriers are among the most extensively investigated systems for oral pDNA delivery in fish^5,11–13^. In gilthead sea bream (*Sparus aurata*), chitosan-encapsulated pDNA enabled reporter gene expression in the intestine, liver, and muscle following oral administration, with detectable expression persisting for up to 60 days^5,11^. Nevertheless, the performance of chitosan-based systems is strongly influenced by polymer characteristics, particularly the degree of deacetylation, molecular weight, and crosslinking conditions^14,15^. Furthermore, the electrostatic interactions between chitosan and DNA may be weakened under the relatively alkaline intestinal conditions of many fish species, potentially leading to premature DNA release and degradation before cellular uptake^16–18^. PLGA nanoparticles also offer favorable biocompatibility and controlled-release properties, although transgene expression and protective immune responses induced by their application in oral pDNA delivery have varied considerably in rainbow trout (*Oncorhynchus mykiss*) and Japanese flounder (*Paralichthys olivaceus*)^12,13,19,20^. Both approaches continue to face challenges in overcoming intestinal barriers and achieving efficient delivery to extraintestinal organs. These limitations highlight the need for alternative carriers with improved biological transport capabilities.

Bovine milk-derived extracellular vesicles (MEVs) have emerged as promising natural carriers for oral drug delivery^21,22^. These nanosized, lipid bilayer-enclosed vesicles exhibit considerable resistance to gastrointestinal degradation, thereby protecting their encapsulated cargo^22–24^. More importantly, studies in mammalian models have demonstrated that MEVs can undergo intestinal epithelial uptake and transepithelial transport, with the neonatal Fc receptor (FcRn) implicated in this process through interactions with vesicle-associated immunoglobulin G (IgG)^25–27^. Such biologically mediated transport may offer an advantage over conventional synthetic carriers that primarily provide physical protection. Although whether similar transport mechanisms operate in fish remains unclear, the gastrointestinal stability and potential intestinal transport capabilities of MEVs make them attractive candidates for oral gene delivery. However, their ability to deliver nucleic acids to extraintestinal tissues, particularly the hepatopancreas, has yet to be established in fish.

Beyond intestinal absorption, successful pDNA delivery also requires efficient DNA condensation, protection, and intracellular trafficking^28,29^. Polyethylenimine (PEI), a widely used cationic polymer for non-viral gene delivery, can electrostatically condense pDNA into nanoscale complexes, enhance its resistance to nuclease degradation, and potentially facilitate endosomal escape^28–31^. However, its dose-dependent cytotoxicity and limited intestinal transport efficiency constrain its application in oral delivery^26,32^. Combining PEI with MEVs may therefore integrate the DNA-condensing and intracellular delivery capabilities of PEI with the gastrointestinal stability and epithelial transport potential of MEVs^33^. Despite these promising properties, the application of MEVs in fish remains largely unexplored. The limited evidence currently available comes primarily from zebrafish (*Danio rerio*), in which the developmental and acute toxicity of bovine milk-derived nanovesicles has been investigated, providing preliminary insights into their biological compatibility in fish^34^. However, these studies did not evaluate oral gene delivery efficiency or transgene expression in target organs of aquaculture species. To our knowledge, successful hepatic transgene expression following oral administration of MEV-based pDNA formulations has not yet been reported in teleosts. We therefore hypothesize that integrating the gastrointestinal stability and potential biological transport properties of MEVs with the DNA-condensing and intracellular delivery capabilities of PEI may provide a promising strategy for non-invasive gene delivery to the fish hepatopancreas.

Building on this rationale, we developed a ternary nanocarrier composed of MEVs, PEI and pDNA, in which PEI was introduced to facilitate DNA condensation and electrostatic complexation. This study aimed to investigate whether the MEV/PEI/pDNA system could enable oral delivery of exogenous pDNA and functional transgene expression in the goldfish hepatopancreas. We first characterized the physicochemical properties, DNA loading capacity, and nuclease protection of the complexes, followed by formulation optimization and evaluation of transfection efficiency and cytocompatibility *in vitro*. The optimized formulation was subsequently incorporated into feed and administered to goldfish over consecutive days to assess the temporal profiles of hepatic reporter gene transcription and protein expression. This study seeks to establish the feasibility of a non-invasive, potentially scalable strategy for feed-based gene delivery in aquaculture and provide a foundation for further investigation of MEV-mediated transport in fish.

## RESULTS

### Preparation and characterization of MPPs

To investigate the potential of MEVs as carriers for feed-based pDNA delivery in fish, we constructed ternary MEV–PEI–pDNA complexes (MPPs) by associating MEVs with the cationic polymer PEI, followed by electrostatic complexation with pDNA (Fig. 1a). Dynamic light scattering (DLS) analysis revealed that free circular pDNA exhibited an apparent hydrodynamic diameter of 462.4 ± 45.7 nm with a relatively broad size distribution (polydispersity index [PDI], 0.37 ± 0.15) (Fig. 1b–d). Following complexation with PEI, the particle size decreased markedly to 69.8 ± 11.0 nm, accompanied by a narrower size distribution (PDI, 0.14 ± 0.06). The resulting MPPs exhibited a hydrodynamic diameter of 120.8 ± 12.0 nm and a PDI of 0.15 ± 0.04. Consistent with these changes, zeta potential measurements showed that the negative surface charge of free pDNA (−21.9 ± 4.1 mV) shifted to a positive value following PEI complexation (18.8 ± 6.6 mV), whereas MPPs exhibited a slightly negative zeta potential of −9.1 ± 3.5 mV (Fig. 1e). Western blot analysis further confirmed the presence of the EV-associated markers CD81 and TSG101 in MPPs, indicating that these characteristic MEV proteins were retained following complex formation (Fig. 1f).

**Figure 1.**
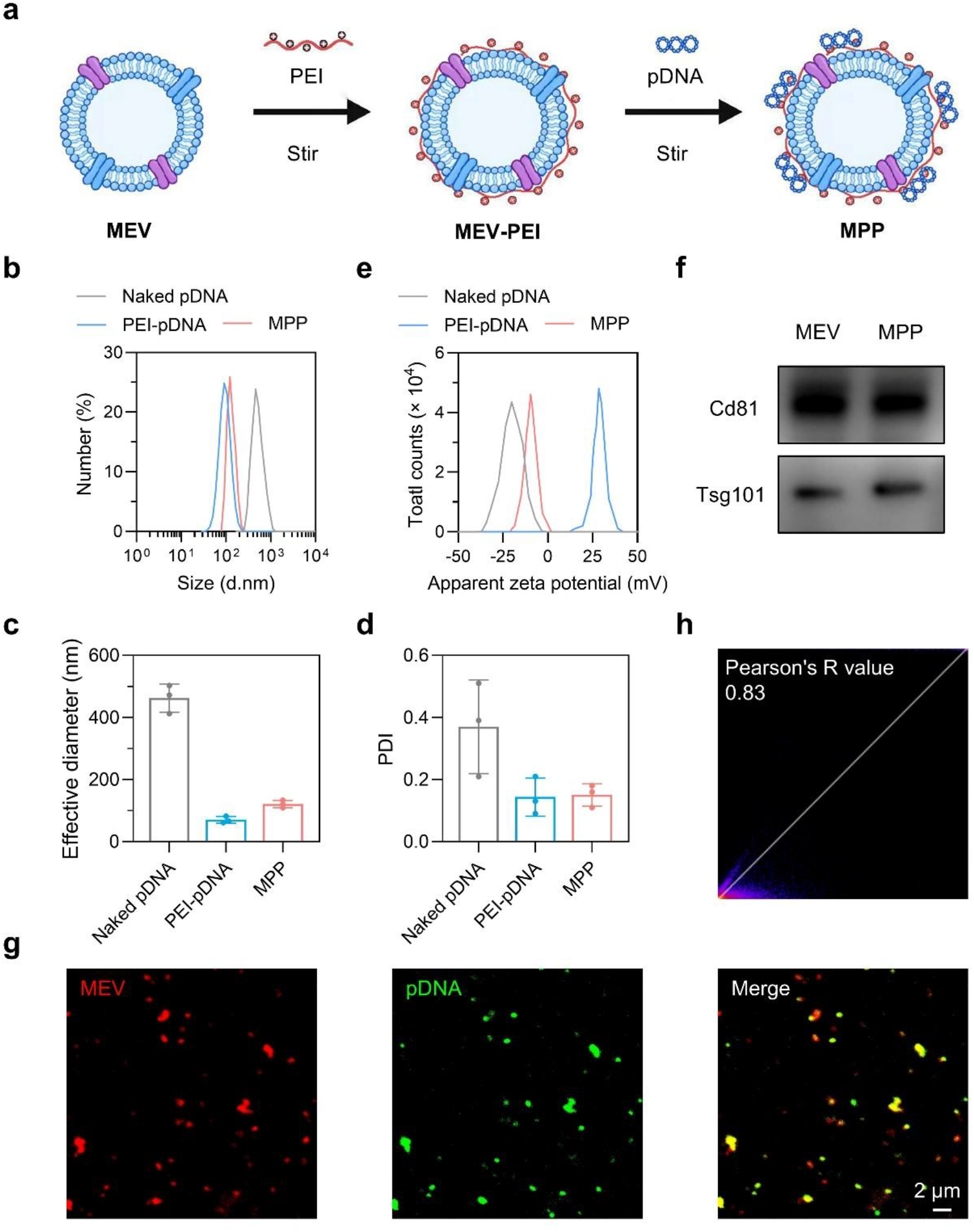
Assembly and physicochemical characterization of MEV–PEI–pDNA complexes. (a) Schematic illustration of MPP preparation. MEV, milk-derived extracellular vesicle; PEI, polyethylenimine; MPP, MEV–PEI–pDNA complex. (b–d) Representative particle size distributions (b), mean hydrodynamic diameters (c), and polydispersity indices (PDIs) (d) of the indicated formulations determined by dynamic light scattering (DLS) (*n* = 3). (e) Zeta potentials of the indicated formulations. (f) Western blot analysis of the EV-associated markers CD81 and TSG101 in MEVs and MPPs. (g) Confocal fluorescence images showing the colocalization of MEVs (red, DiD) and pDNA (green, YOYO-1) in MPPs. (h) Quantitative colocalization analysis of MEV and pDNA fluorescence signals. Data are presented as mean ± SD.

To further examine the association between MEVs and pDNA, MEVs were labeled with the lipophilic fluorescent dye DiD, while pDNA was labeled with the DNA-intercalating dye YOYO-1 prior to complex assembly. Confocal fluorescence microscopy revealed substantial spatial overlap between the two fluorescent signals (Fig. 1g), which was further supported by colocalization analysis showing a Pearson’s correlation coefficient of 0.83 (Fig. 1h). Finally, pDNA loading was quantified using YOYO-1-labeled pDNA by measuring the amount of unbound fluorescent pDNA remaining after complex formation. The resulting MPPs exhibited a pDNA loading efficiency of 98.5 ± 2.4% and a loading content of 3.9 ± 1.1%. Collectively, these results demonstrate the successful formation of MPPs with well-defined physicochemical characteristics, high pDNA loading efficiency, and close association between the vesicular and DNA components, providing a basis for subsequent evaluation of their gene delivery performance.

### Payload release and stability of MPPs

The stability and release behavior of gene delivery carriers are important determinants of their performance following oral administration. We therefore first investigated the *in vitro* pDNA release kinetics of MPPs under simulated physiological conditions (Fig. 2a). MPPs exhibited a sustained release profile, with an initial release of approximately 25% of the loaded pDNA within the first 2 h, followed by a slower increase to approximately 63% at 24 h. Cumulative release approached 90% and reached a plateau between 48 and 72 h. This biphasic release pattern may reflect the relatively rapid dissociation of loosely associated pDNA, followed by the gradual release of DNA more strongly retained within the complexes. We next evaluated the ability of MPPs to protect pDNA against enzymatic degradation using a DNase I protection assay. Agarose gel electrophoresis showed no detectable pDNA band following DNase I treatment of either free pDNA or MPPs (Fig. 2b). Given that PEI-mediated DNA condensation can restrict electrophoretic migration and interfere with fluorescent dye binding, sodium heparin was added to competitively displace pDNA from PEI^30^. Following heparin treatment, a distinct pDNA band was recovered from DNase I-treated MPPs, indicating that complexation provided protection against nuclease-mediated degradation. Such protection is particularly relevant to oral gene delivery, where nucleic acids are exposed to degradative conditions in the gastrointestinal tract. Finally, we assessed the storage stability of MPPs by monitoring their hydrodynamic diameter at 4 °C. Particle size remained relatively stable during the first 3 days but increased to approximately 250 nm after 7 days (Fig. 2c), suggesting that prolonged storage may compromise their colloidal stability. These findings demonstrate that MPPs provide sustained pDNA release and nuclease protection, while highlighting the importance of appropriate storage conditions for their subsequent application in feed-based gene delivery.

**Figure 2.**
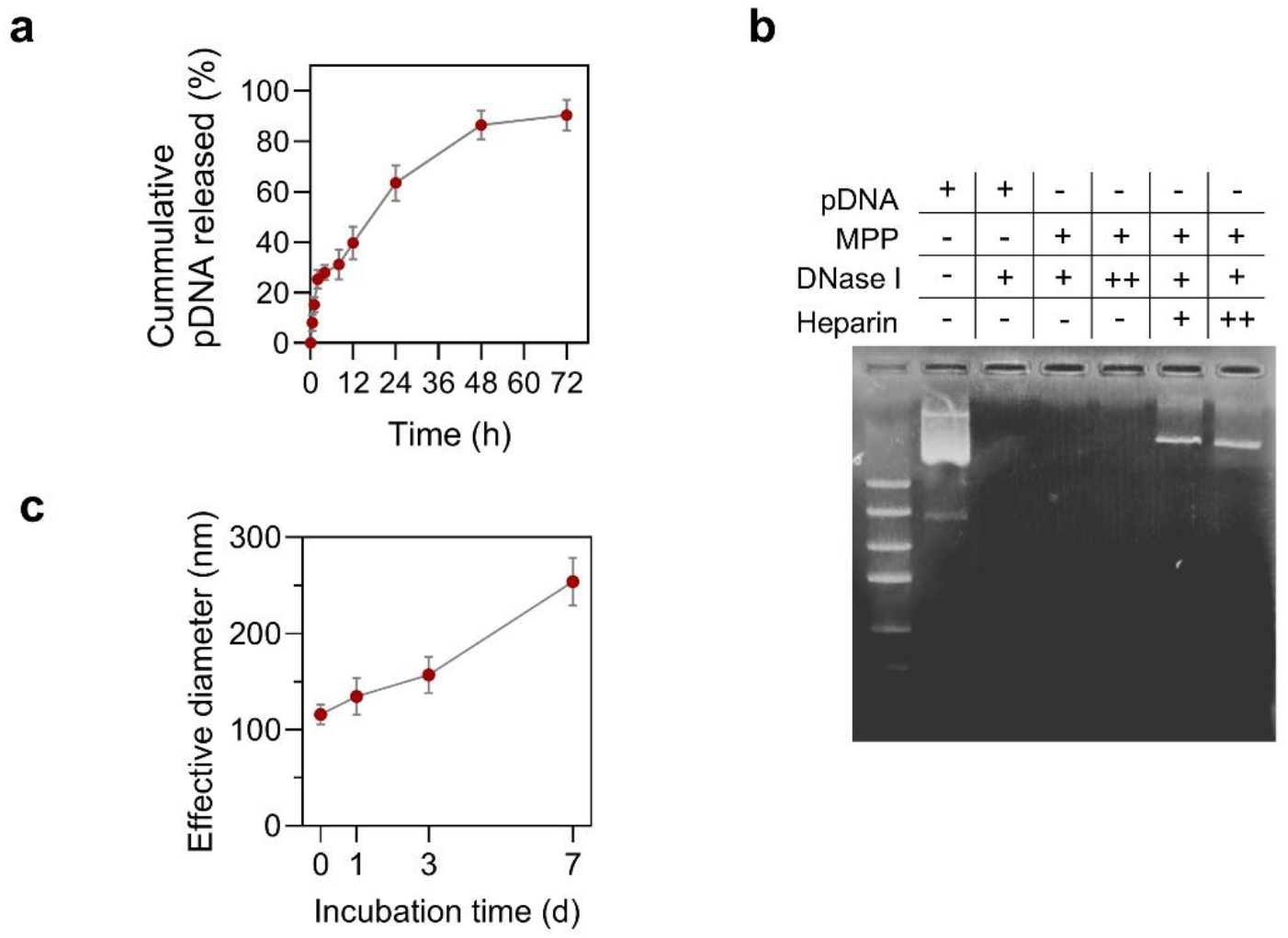
Release behavior, nuclease protection, and storage stability of MPPs. (a) *In vitro* pDNA release profile of MPPs under simulated physiological conditions (*n* = 3). (b) Agarose gel electrophoresis analysis of the protective effect of MPPs against DNase I-mediated pDNA degradation. Sodium heparin was used to competitively displace pDNA from PEI. For DNase I, “+” and “++” indicate concentrations of 10 and 20 U/mL, respectively; for heparin, “+” and “++” indicate concentrations of 200 and 400 μg/mL, respectively. (c) Changes in the hydrodynamic diameter of MPPs during storage at 4 °C (*n* = 3). Data are presented as mean ± SD.

### Optimization of MPP formulation for *in vitro* transfection

The transfection performance of MPPs is influenced by the relative proportions of their three components. In particular, the nitrogen-to-phosphate (N/P) ratio of PEI to DNA can affect transfection efficiency by modulating DNA condensation, nuclease protection, endosomal escape, and cytotoxicity^35^, whereas the MEV-to-pDNA ratio may influence the physicochemical properties and cellular interactions of the resulting complexes^33^。We first investigated the effect of PEI content on MPP-mediated transfection using EGFP as a reporter gene in HEK293T cells. With the amounts of MEVs and pDNA fixed at 37.5 and 2.25 μg per well, respectively, in 48-well plates containing 250 μL of culture medium per well, the PEI content was varied relative to the pDNA mass. Fluorescence microscopy revealed a progressive increase in EGFP expression with increasing PEI content, reaching the highest level at a PEI mass ratio of 4.1:1 (9.25 μg PEI per well), with fluorescence signals comparable to those obtained using Lipofectamine 3000. Further increases in PEI content did not improve EGFP expression, while a marked reduction in cell density was observed at a PEI mass ratio of 16.4:1, suggesting dose-dependent cytotoxicity (Fig. 3a). Accordingly, a PEI:pDNA mass ratio of 4.1:1 was selected for subsequent optimization.

**Figure 3.**
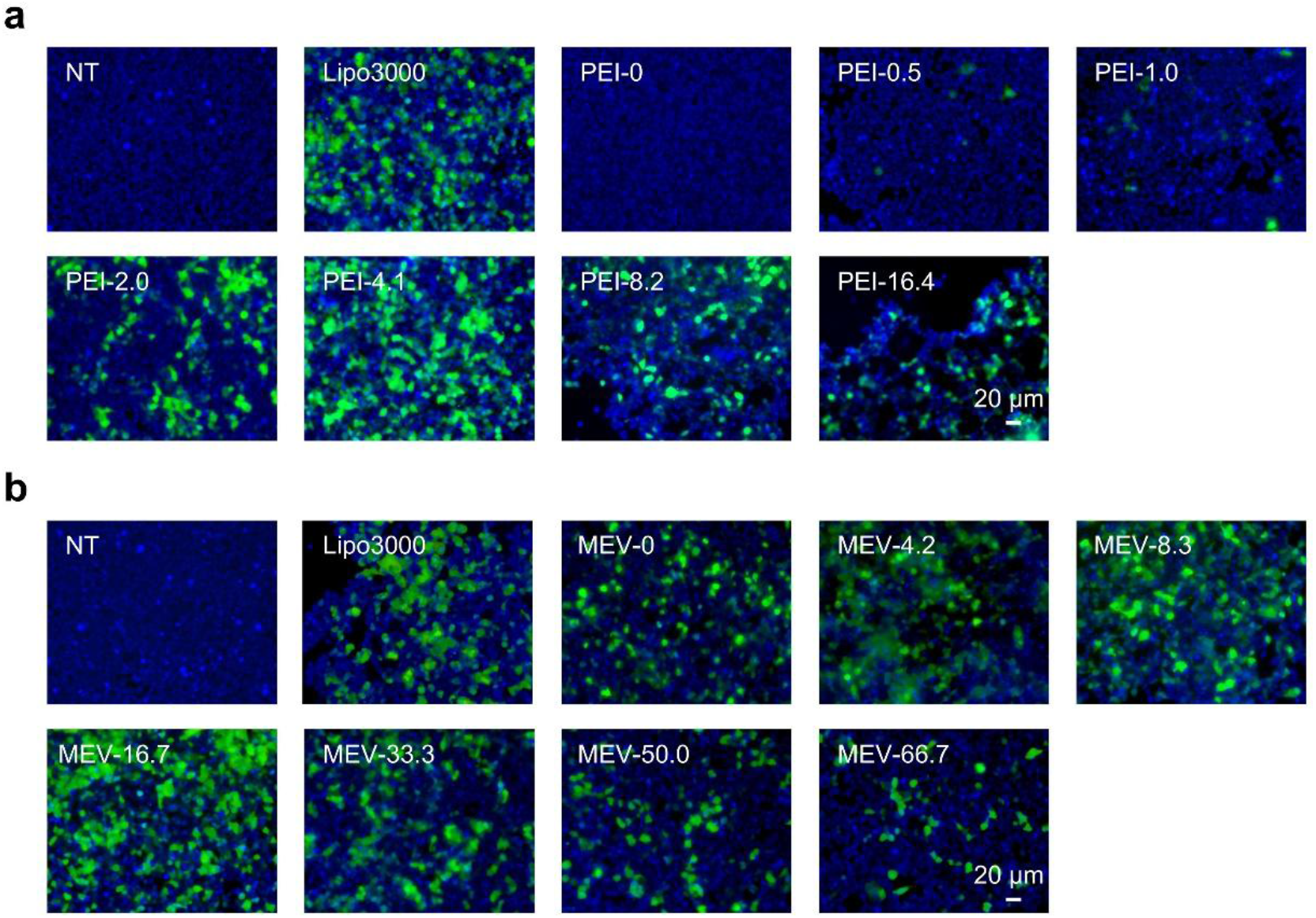
Effect of PEI and MEV content on MPP-mediated EGFP expression. (a) The formulation of MPPs was optimized by varying the mass ratios of PEI or MEV to pDNA, using enhanced green fluorescent protein (EGFP) as a reporter in HEK293T cells cultured in 48-well plates. (a) Representative fluorescence micrographs showing EGFP expression following transfection with MPPs prepared at different PEI:pDNA mass ratios, with the amounts of MEVs and pDNA fixed at 37.5 and 2.25 μg per well, respectively. (b) Representative fluorescence micrographs showing EGFP expression following transfection with MPPs prepared at different MEV:pDNA mass ratios, with the amounts of PEI and pDNA fixed at 9.25 and 2.25 μg per well, respectively. Untreated HEK293T cells (NT) served as the negative control, while cells transfected using the commercial reagent Lipofectamine 3000 served as the positive control. Scale bar, 20 μm; Green, EGFP; blue, cell nuclei.

We next examined the effect of MEV content by maintaining the amounts of PEI and pDNA at 9.25 and 2.25 μg per well, respectively, while varying the MEV mass ratio. No apparent cytotoxic effects were observed across the tested MEV concentrations. However, EGFP expression initially increased with increasing MEV content and subsequently declined, reaching its highest level at an MEV:pDNA mass ratio of 16.7:1 (Fig. 3b). This non-monotonic response suggests that an appropriate balance between MEVs and PEI as well as pDNA is important for efficient transfection, as excessive MEV incorporation may alter complex assembly or cellular interactions. Together, these results indicate that both PEI and MEV contents influence MPP-mediated gene expression, with an optimal balance required to achieve efficient transfection while minimizing potential cytotoxicity. Based on these findings, an MEV:PEI:pDNA mass ratio of 16.7:4.1:1 was selected for subsequent experiments.

### Feed-based delivery and hepatic transgene expression of MPPs in goldfish

Following the *in vitro* evaluation of transfection efficiency and cytocompatibility, we next investigated whether MPPs could deliver exogenous pDNA to the goldfish liver (hepatopancreas) and achieve functional transgene expression following oral administration through feed. Firefly luciferase (Luc) was selected as the reporter gene, and MPPs were thoroughly mixed with fish feed before administration to acclimated goldfish. To assess the tissue distribution of MPPs, the MEV component was labeled with the lipophilic fluorescent dye DiD, and major organs were collected at different time points after feeding for *ex vivo* fluorescence imaging. IVIS imaging revealed detectable background fluorescence in the intestinal tract before feeding (0 h), despite a 24-h fasting period, potentially reflecting residual intestinal contents or tissue autofluorescence (Fig. 4a). At 2 h post-feeding, intestinal fluorescence increased markedly, and a detectable signal also appeared in the hepatopancreas, whereas no appreciable fluorescence was observed in the other examined organs. The hepatopancreatic signal remained detectable at 6 h but was no longer apparent at 24 h, when a weak fluorescent signal emerged in the kidney. These observations suggest that MEV-associated fluorescent components can reach the hepatopancreas following oral administration and subsequently undergo redistribution or probably renal excretion.

**Figure. 4.**
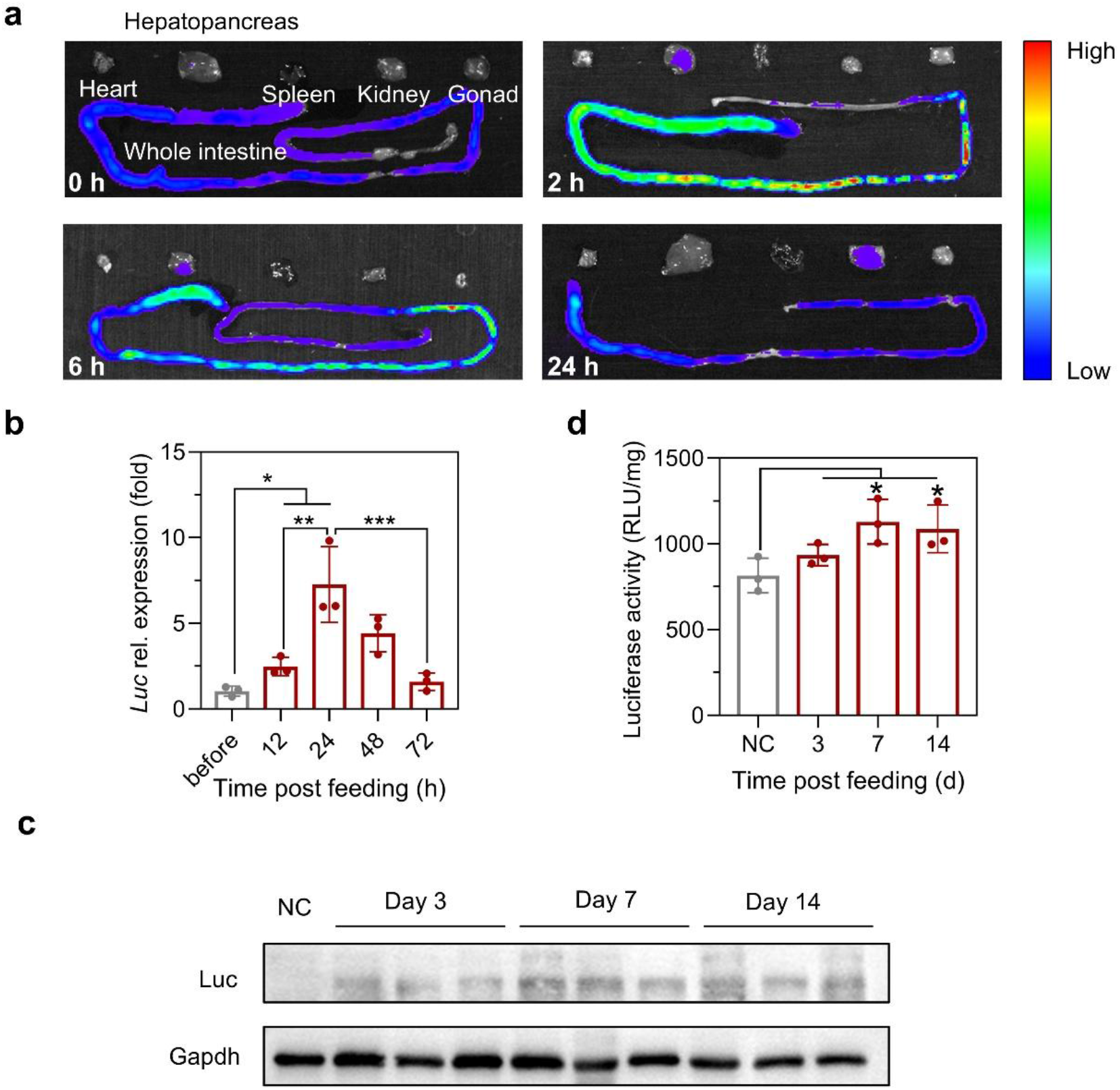
Tissue distribution and functional transgene expression following feed-based administration in goldfish. (a) *Ex vivo* fluorescence imaging of major organs collected from goldfish at the indicated time points following a single feeding of DiD-labeled MEVs incorporated into fish feed. Fluorescence signals were detected using an *in vivo* imaging system (IVIS). (b) Relative luciferase mRNA expression in the hepatopancreas at different time points following a single feeding of MPP-containing feed, as determined by RT-qPCR (*n* = 3). The *gapdh* gene was used as the internal reference for normalization. (c) Western blot analysis of luciferase protein expression in the hepatopancreas at the indicated time points during 14 consecutive days of feeding with MPP-containing feed. (d) Luciferase activity in hepatopancreatic tissue homogenates at the indicated time points during the 14-day feeding period (*n* = 3). Luciferase activity was normalized to tissue weight and expressed as relative light units per milligram of tissue (RLU/mg tissue). Data are presented as mean ± SD. \**P* < 0.05, \*\**P* < 0.01, and \*\*\**P* < 0.001.

To determine whether this tissue distribution was accompanied by functional gene delivery, we first examined hepatic *Luc* mRNA expression following a single feeding. RT-qPCR analysis revealed that Luc transcript levels peaked at 24 h post-feeding and subsequently declined, approaching baseline levels by 72 h (Fig. 4b). We then investigated whether repeated administration could support sustained transgene expression by feeding goldfish an MPP-containing diet daily for 14 consecutive days. Western blot analysis detected weak Luc protein expression on day 3, followed by progressively stronger signals on days 7 and 14 (Fig. 4c). Consistent with these findings, luciferase activity measured in hepatopancreatic tissue homogenates exhibited a similar temporal pattern, with higher enzymatic activity observed on days 7 and 14 (Fig. 4d). The transient transcriptional response following a single feeding, together with the increased reporter protein expression and enzymatic activity during repeated administration, suggests that continued dietary delivery may help maintain functional transgene expression. Collectively, these findings demonstrate that feed-based administration of MPPs enables detectable exogenous gene expression in the goldfish hepatopancreas, supporting the feasibility of this non-invasive gene delivery strategy for aquaculture applications.

## CONCLUSION

In conclusion, this study provides proof of concept for the use of MEVs as carriers for feed-based pDNA delivery in fish. We developed a ternary nanocomplex comprising MEVs, PEI, and pDNA (MPP) and systematically evaluated its physicochemical properties, DNA loading and protection, *in vitro* transfection performance, and oral delivery potential. MPPs exhibited a relatively uniform nanoscale size distribution, high pDNA loading efficiency, sustained release, and protection against nuclease degradation. Following formulation optimization, MPPs achieved robust transgene expression *in vitro* with favorable apparent cell tolerance. More importantly, dietary administration of MPPs to goldfish resulted in detectable MEV-associated fluorescence in the hepatopancreas, accompanied by reporter gene transcription, protein expression, and luciferase activity. The increased reporter activity observed during repeated feeding further suggests that continued oral administration may help maintain functional transgene expression. Together, these findings support the feasibility of MEV/PEI-based oral gene delivery and provide a foundation for developing non-invasive, scalable genetic delivery strategies in aquaculture. Nevertheless, further investigations are needed to elucidate the intestinal transport mechanisms and tissue distribution of intact MPPs, establish dose–response relationships and long-term expression profiles, and address potential toxicity, immunogenicity, formulation stability, and production scalability before practical aquaculture applications can be considered.

## METHODS

### Ethics statement

All animal experiments were conducted in accordance with the *Guide for the Care and Use of Laboratory Animals* published by the National Research Council and were approved by the Institutional Animal Care and Use Committee (IACUC) of China Science Industry Holding (Shenzhen) Co., Ltd. (approval No. 20260018).

### Cell lines and animals

Human embryonic kidney 293T (HEK293T) cells were maintained in Dulbecco’s modified Eagle’s medium (DMEM; Gibco) supplemented with 10% heat-inactivated fetal bovine serum (FBS; Gibco), 100 U/mL penicillin, and 100 μg/mL streptomycin at 37 °C in a humidified atmosphere containing 5% CO₂.

Goldfish (*Carassius auratus*) weighing 250–300 mg were obtained from local suppliers and individually housed in 2-L aquaria at 20–25 °C under a 12-h light/12-h dark photoperiod. Fish were acclimated to laboratory conditions for at least 2 weeks before experimentation. Prior to tissue collection, goldfish were immersed in a solution containing 1 g/L tricaine methanesulfonate (MS-222; Sigma-Aldrich) for terminal anesthesia.

### Isolation of milk-derived extracellular vesicles

Bovine milk-derived extracellular vesicles (MEVs) were isolated from freshly collected raw bovine milk according to a previously reported method with minor modifications^36^。Briefly, fresh milk was centrifuged at 2,500 × g for 30 min at 4 °C to remove the cream layer, somatic cells, and cellular debris. The resulting defatted milk was supplemented with ethylenediaminetetraacetic acid (EDTA) to a final concentration of 30 mM and incubated at 37 °C for 1 h to disrupt calcium-dependent casein micelles. The mixture was subsequently subjected to sequential centrifugation at 2,500 × g and 12,000 × g for 1 h each, with the supernatant collected after each step. The resulting supernatant was passed through a 0.22-μm syringe filter and ultracentrifuged at 140,000 × g for 1 h to obtain an MEV-enriched pellet. The pellet was resuspended in phosphate-buffered saline (PBS) and washed by an additional ultracentrifugation step at 100,000 × g for 1 h at 4 °C. Finally, the MEV pellet was resuspended in PBS and stored at −80 °C until further use.

### Preparation of the MPPs

MEV–PEI–pDNA complexes (MPPs) were prepared using a previously reported method with minor modifications^33^。Briefly, MEVs were initially mixed with linear polyethylenimine (PEI; 25 kDa; Macklin), followed by the addition of plasmid DNA (pDNA) to facilitate electrostatic complex formation. In a typical reaction, MEVs, PEI, and pDNA were combined at a mass ratio of 16.7:4.1:1. One volume of PEI solution (2 μg/μL) was mixed with an equal volume of MEV suspension by vortexing, followed by incubation at room temperature for 5 min. Subsequently, two volumes of pDNA solution were added, and the mixture was vortexed and incubated at room temperature for an additional 15 min. The resulting MPPs were processed using centrifugal ultrafiltration devices with a molecular weight cutoff (MWCO) of 300 kDa before further analysis.

### Physicochemical characterization and fluorescence labeling of MPPs

The hydrodynamic diameter, polydispersity index (PDI), and zeta potential of MPPs were determined using a Zetasizer instrument (Malvern Panalytical). Particle concentration was measured by nanoparticle tracking analysis (NTA) using a ZetaView analyzer (Particle Metrix) equipped with a Z-NTA cell.

For fluorescence imaging, MEVs and pDNA were separately labeled with the lipophilic membrane dye DiD and the DNA-intercalating dye YOYO-1, respectively, before MPP assembly. To label MEVs, equal volumes of MEV suspension (1 mg/mL) and DiD solution (10 μM) were gently mixed and incubated at 37 °C for 30 min in the dark. Unbound DiD was subsequently removed using a Sephadex G-25 column. For pDNA labeling, equal volumes of pDNA solution (1 mg/mL) and YOYO-1 solution (20 μM; MedChemExpress) were gently mixed and incubated at room temperature for 30 min. Unbound YOYO-1 was then removed using a Sephadex G-25 column. The fluorescently labeled MEVs and pDNA were subsequently used to prepare MPPs as described above. The resulting fluorescent complexes were directly loaded on glass slides, covered with coverslips, and examined using an LSM 710 confocal laser scanning microscope (Carl Zeiss).

### Western blot analysis

Western blotting was performed to detect the EV-associated proteins CD81 and TSG101 in MEV preparations and MPPs, as well as luciferase (Luc) expression in goldfish hepatopancreatic tissues. Total protein was extracted using a Mammalian Total Protein Extraction Kit (TransGen Biotech) supplemented with protease and phosphatase inhibitor cocktails (Thermo Fisher Scientific). Protein samples were separated by sodium dodecyl sulfate–polyacrylamide gel electrophoresis (SDS-PAGE) and transferred onto polyvinylidene fluoride (PVDF) membranes at a constant current of 150 mA for 1.5 h. The membranes were blocked with 5% bovine serum albumin (BSA) for 1.5 h at room temperature and subsequently incubated overnight at 4 °C with primary antibodies against CD81 (1:1,000; Abcam), TSG101 (1:2,000; Abcam), Luc (1:1,000; Cell Signaling Technology), or GAPDH (1:10,000; Abcam). After washing, the membranes were incubated with the appropriate horseradish peroxidase (HRP)-conjugated anti-rabbit or anti-mouse IgG (H+L) secondary antibodies (1:2,000; Cell Signaling Technology) for 1 h at room temperature. Immunoreactive bands were detected using an enhanced chemiluminescence (ECL) substrate (Thermo Fisher Scientific) and visualized using an Amersham Imager 600 imaging system (GE Healthcare).

### pDNA loading and *in vitro* release

The pDNA loading content and loading efficiency of MPPs were evaluated using YOYO-1-labeled pDNA. Briefly, fluorescently labeled pDNA was incorporated into MPPs during complex preparation as described above. The resulting formulations were subjected to centrifugal ultrafiltration using 300-kDa molecular weight cutoff (MWCO) filters, and the fluorescence intensity of the filtrate was measured at excitation and emission wavelengths of 491 and 509 nm, respectively, to estimate the amount of unbound pDNA. The pDNA loading content (LC) and loading efficiency (LE) were calculated using the following equations:

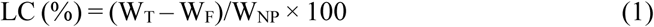

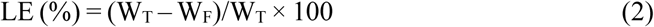

where W_T_ represents the total mass of pDNA initially added, W_F_ represents the estimated mass of unbound pDNA and W_NP_ represents the mass of MPPs.

The *in vitro* release profile of pDNA from MPPs was evaluated using a dialysis-based method with modifications^37^. MPPs containing YOYO-1-labeled pDNA were placed in dialysis membranes (MWCO, 300 kDa) and immersed in PBS supplemented with 5% FBS at 37 °C under continuous agitation. At predetermined time points, aliquots of the surrounding release medium were collected for fluorescence measurement and replaced with equal volumes of fresh medium.

### DNase I protection assay

The ability of MPPs to protect pDNA against nuclease-mediated degradation was evaluated using a DNase I protection assay. Free pDNA (1.25 μg) or MPPs containing an equivalent amount of pDNA were incubated with DNase I (10 or 20 U/mL; Yeasen) at 37 °C for 10 min. The reaction was terminated by adding EDTA to a final concentration of 5 mM, followed by incubation at 80 °C for 20 min. For MPP samples, an equal volume of sodium heparin solution (200 or 400 μg/mL in 300 mM NaCl) was subsequently added to competitively displace pDNA from PEI^30^. The mixtures were vigorously agitated at room temperature for 15 min and centrifuged at 12,800 × g for 2 min. The resulting supernatants containing released pDNA were collected and analyzed by electrophoresis on 1.5% agarose gels at 135 V for 30 min.

### *In vitro* gene transfection

The *in vitro* transfection performance of MPPs was evaluated in HEK293T cells using enhanced green fluorescent protein (EGFP) as a reporter gene. Cells were seeded into 48-well plates at a density of 5 × 10⁴ cells per well and cultured overnight until approximately 70–80% confluence was reached. The culture medium was then replaced with fresh serum-free medium, and cells were incubated with MPPs prepared with varying PEI:pDNA or MEV:pDNA mass ratios. After 24 h, the transfection medium was replaced with fresh complete growth medium, and the cells were cultured for an additional 24 h. EGFP fluorescence and cellular morphology were subsequently examined using an inverted fluorescence microscope (CKX53; Olympus).

### Goldfish feeding experiment

Goldfish were individually maintained in 2-L aquaria and acclimated to a regular feeding schedule before experimentation. During the acclimation period, fish were conditioned to consume the entire feed ration within 2 min of feed introduction. Each fish was provided with 24 mg of commercial fish feed per feeding, twice daily at 10:00 and 16:00.

To prepare the MPP-containing feed, 12 μL of MPP suspension containing 1.12 μg of pDNA was added dropwise to 24 mg of fish feed under vigorous mixing to ensure uniform distribution of the formulation. The prepared feed was administered immediately after mixing. Throughout the feeding experiment, goldfish received the MPP-containing feed twice daily at the designated times, and samples were collected at predetermined time points for subsequent analyses.

### *In vivo* tissue distribution

To investigate the tissue distribution of orally administered MEVs, DiD-labeled MEVs were mixed with 24 mg of fish feed and administered to goldfish as a single feeding. At predetermined time points after feeding, fish were euthanized, and major organs were collected for *ex vivo* fluorescence imaging using an IVIS Lumina III imaging system (PerkinElmer).

### Quantitative analysis of Luc mRNA expression

To evaluate reporter gene transcription following oral administration, goldfish received a single feeding of 24 mg of MPP-containing feed. At predetermined time points after feeding, fish were euthanized, and hepatopancreatic tissues were immediately collected and homogenized in TRIzol reagent. Total RNA was extracted according to the manufacturer’s instructions and reverse-transcribed into complementary DNA (cDNA) using the RevertAid First Strand cDNA Synthesis Kit (Thermo Fisher Scientific). Quantitative real-time PCR (qPCR) was performed using a CFX96 Connect Real-Time PCR Detection System (Bio-Rad) with Real-Time SYBR Green I PCR Master Mix (Yeasen). The amplification conditions consisted of an initial denaturation at 95 °C for 5 min, followed by 40 cycles of denaturation at 95 °C for 10 s, annealing at 60 °C for 20 s, and extension at 72 °C for 20 s.

Primer sequences were as follows: Luc-F: 5’-ACTGGGACGAAGACGAACAC-3’; Luc-R: 5’-TGCTCCAAAACAACAACGGC-3’; gapdh-F: 5’-GATACACGGAGGACCAGGTG-3’; gapdh-R: 5’-TGGGTCGTTTACTCCTTGGAT-3’.

### Luciferase activity assay

At predetermined time points during the feeding experiment, goldfish were euthanized, and hepatopancreatic tissues were collected, weighed and mechanically homogenized. Luciferase activity was determined using a Luciferase Assay System kit (Promega). The homogenized tissue samples were processed for the assay, and luminescence was measured using a Varioskan Flash multimode microplate reader (Thermo Scientific) with a signal integration time of 10 s. Luciferase activity was normalized to the weight of the corresponding hepatopancreatic tissue and expressed as relative light units per milligram of tissue (RLU/mg tissue).

### Statistical analysis

Statistical analyses were performed using GraphPad Prism 9.0 software. Differences among multiple groups were evaluated by one-way analysis of variance (ANOVA), followed by Tukey’s, Dunnett’s, or Bonferroni-adjusted multiple-comparison tests, as appropriate. A *P* value of less than 0.05 was considered statistically significant.

## Acknowledgments

This work was supported by grants from the Shenzhen-Hong Kong Institute of Brain Science-Shenzhen Fundamental Research Institutions (2023SHIBS0003).

## Author contributions

B.Z. and S.L. designed the study. B.Z. and S.L. analyzed the data. B.Z. prepared the manuscript. B.Z. and S.L. reviewed and edited the manuscript. Q.Z., X.M. and W.L. prepared and characterized the nanoparticles. Q.Z., and X.M. carried out cell experiments. Q.Z., and X.M. conducted animal experiments.

## Competing interests

The authors declare no competing financial interests.

## Notes

### Competing Interest Statement

The authors have declared no competing interest.

